# A Modular Bio-Hybrid Skin Model for Optical Testing Applications

**DOI:** 10.64898/2026.07.27.740463

**Authors:** Dardan Bajrami, Kongchang Wei, Fabrizio Spano, Nolan Agah, Mathias Bonmarin, René M. Rossi

**Affiliations:** Empa, Swiss Federal Laboratories for Materials Science and Technology, Laboratory for Biomimetic Membranes and Textiles, Lerchenfeldstrasse 5, St.Gallen 9014, Switzerland; Zurich University of Applied Sciences (ZHAW), School of Engineering, Technikumstrasse 71, Winterthur 8401, Switzerland; ETH Zürich, Department of Health Science and Technology, Zürich, Switzerland; Empa, Swiss Federal Laboratories for Materials Science and Technology, Laboratory for Biointerfaces, Lerchenfeldstrasse 5, St. Gallen 9014, Switzerland

**Keywords:** Optical model, Light–tissue interaction, Bio-hybrid skin model, skin model

## Abstract

Synthetic optical skin models offer reproducible, tunable optical properties but lack biological responsiveness, while tissue engineered skin models provide cellular authenticity but suffer from optical variability and limited controllability. The growing demand for alternatives to animal models in the development and validation of optical biomedical technologies highlights the need for a new class of test system that combines the strengths of both approaches while addressing their respective limitations. Here, we introduce the concept of a modular biohybrid skin model, a new testing concept that integrates an optically defined artificial epidermal layer, incorporating polydopamine nanoparticles for changes in skin tone, with living human keratinocytes in two and three-dimensional configurations. In the Optical Protection Model, UV-B-induced apoptosis in primary keratinocytes is quantitatively modulated by model pigmentation level, demonstrating a relationship between optical attenuation and caspase 3/7 activity across three artificial skin tone conditions. In a Structured Dermal Model, keratinocytes seeded onto a hydrogel scaffold localize within follicle-like microcavities, as confirmed by live/dead staining and confocal z-stack imaging. Together, these experiments lead to a new category of test system in the space between inert optical models and variable tissue models that may contribute to reducing the reliance on animal models in biomedical optics.

## Introduction

The increasing integration of optical technologies into biomedical devices, diagnostic systems, and wearables is leading to a growing need for reliable and reproducible test platforms for the systematic investigation of light-tissue interactions ^1, 2^. The efficacy and safety of a wide range of light-based biomedical applications are determined by the optical and biological properties of the skin ^3–5^. However, current strategies are often based either on purely optical models ^6–11^ with limited biological relevance or on *ex vivo* and *in vitro* models that are associated with high variability, limited controllability, and ethical limitations ^12–15^. Classical optical skin models are typically based on a matrix material such as silicone elastomers or polymer hydrogel, with embedded light scattering and absorbing particles, and offer excellent reproducibility and precisely tunable optical properties ^1, 16^. The systems have proven invaluable for calibration, benchmarking, and initial device characterization. However, their purely synthetic nature limits their similarity to real skin by not being able to capture cellular response coming from optical stimuli such as photodamage or biochemical signaling ^2, 17^. This constraint becomes particularly limiting when developing technologies intended to interact with living tissue, where the biological response is as critical as the optical interaction itself, such as hair follicle removal, sunscreen testing, and photodynamic therapy.

Conversely, tissue-engineered *in vitro* skin models and *ex vivo* tissue samples provide biological authenticity but suffer from inherent variability in optical properties. Donor-dependent factors, including melanin content, collagen density, vascularization, and hydration state, introduce substantial inter-sample variation ^3, 18–20^. *Ex vivo* models further compound this through experimental variables such as tissue freshness, preparation protocols, and sample thickness ^21^. Such models remain valuable for late-stage validation, yet the inherent optical variability limits their utility in early-stage, iterative development workflows where controlled experimental conditions are required. The variance in scattering and absorption characteristics can confound experimental outcomes and complicate the interpretation of results gathered on biological material. Moreover, ethical considerations, high costs, and limited throughput further restrict their use in iterative development cycles ^21, 22^. This mismatch between optically defined models and biologically responsive models is a critical bottleneck in the development of optical biomedical technologies. Ideally, test models should combine controlled, reproducible optical properties that enable systematic studies, living cellular components capable of responding to optical stimuli, and modularity that enables adaptation to different testing scenarios.

Building upon our previously reported artificial epidermis layer, which successfully replicated the optical properties of human skin across different pigmentation levels ^23^, we present here a bio-hybrid model that integrates living cellular components into an optically defined framework. The aim is to provide a modular, reproducible, and optically relevant skin model. This bio-hybrid model is demonstrated using two exemplary configurations: the Optical Protection Model in which keratinocytes are covered by optically tunable artificial epidermal layers, and the Structured Dermal Model, a hydrogel scaffold with follicle-like microcavities populated with keratinocytes (Figure 1)

**Figure 1.**
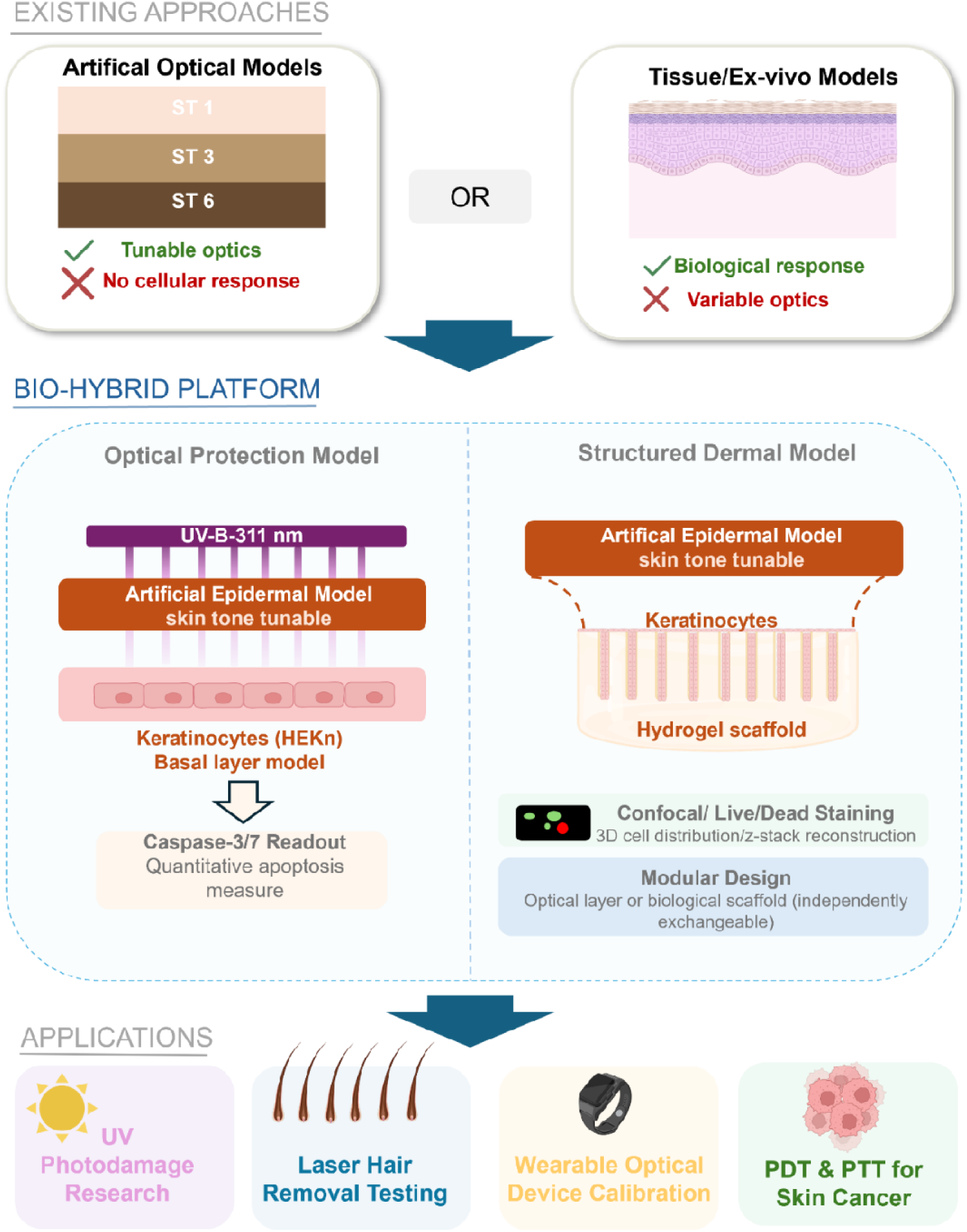
Schematic overview of the bio-hybrid skin model combining a skin-tone-tunable epidermal model with livin keratinocytes in the Optical Protection and Structured Dermal configurations

The usage of this model is clearly exemplified in the context of skin cancer research, where quantitative investigation of UV-induced photodamage across skin tones demands precisely the combination of optical control and biological responsiveness that neither existing model category can provide alone. Skin cancer remains the most diagnosed malignancy worldwide, with its incidence continuing to rise across populations ^24^. Ultraviolet radiation is the dominant risk factor and determines which cell populations receive mutagenic doses ^25–27^. The basal layer of the epidermis is of particular interest, as the proliferating keratinocytes that are the origin of basal cell carcinoma are located in the area ^24, 25, 28^. The UV dose reaching these basal cells is critically dependent on the optical properties of the overlying epidermal layers. A relationship that varies substantially across skin tones due to differences in melanin content and distribution ^3, 29^. Studying these relationships quantitatively requires a model that combines reproducible, skin-tone-specific optical properties with living cells capable of measurable biological responses. In the Optical Protection Model, the living keratinocyte monolayer represents the basal layer of the epidermis. The overlying optical artificial epidermis mimics the remaining stratified layers of the epidermis optically. This abstraction allows the isolation of the biologically most relevant target cells while maintaining controlled, reproducible optical boundary conditions. Beyond the UV irradiation use case, many optical photomedicine applications target structures within the dermis and rely on the biological response of dermal components such as hair follicles in laser hair removal, blood vessels in the treatment of port wine stains ^30^, or photodynamic therapy for dermal tumors. In the Structured Dermal Model, the biohybrid concept illustrates the potential for broader applicability beyond the specific UV protection use case. It demonstrates the model’s capacity to generate geometrically defined, cell-populated structures within an optically characterized framework with different skin colors. By populating a 3D printed hair follicle structure with relevant cell types (keratinocytes), we show how the model evolves into a living optical test system, which could be used for numerous applications.

Collectively, these two demonstrators illustrate the fundamental concept underlying this model: the ability to combine standardized optical properties with biological components in modular configurations. The modular nature of the model enables researchers to select the biological use case that best suits their specific application, whether that involves simple cellular viability assays, functional imaging, or more sophisticated readouts. Similarly, the optical properties can be tuned to absorption levels (skin colors) relevant to specific technologies. This flexibility is intended to inspire and enable diverse applications across photomedicine, wearable sensing, cosmetic testing, and optical device development. The model could therefore offer a new way of testing between traditional optical skin models and tissue-engineered models, and could, in alignment with the 3R principle, contribute to reducing the reliance on animal models ^31^.

## Materials and Methods

### Materials

For the Optical Protection Model: Human epidermal keratinocytes, neonatal (HEKn; Gibco, Thermo Fisher Scientific, Waltham, MA, USA; Catalog #C-001-5C; neonatal male donor, light pigmentation, p3-4. CnT-PR medium (CELLnTEC Advanced Cell Systems, Bern, Switzerland) were used for cell culture. The artificial epidermal layer was derived from the previously reported optical artificial skin ^23^. Staurosporine (Sigma-Aldrich, St. Louis, MO, USA) was used as a positive control for apoptosis induction. Optical irradiation of cells in 2D was performed using a custom-built in-house lamp, including a TL-40W/01 RS UV-B lamp (Philips, Eindhoven, Netherlands) with an emission range of 280–320 nm and a peak wavelength of 311 nm. Apoptosis quantification was performed using Caspase-Glo® assay kit (Promega, Madison, WI, USA). For the Structured Dermal Model: HEKn (neonatal donor, p 3-4) were used for cell seeding. A cold-water fish gelatin based hydrogel (cfGel-Hydrogel) was used as the matrix. The detailed synthesis information was reported by Hammer et al. ^32^. Irgacure 2959 (Sigma-Aldrich, St. Louis, MO, USA) was used for hydrogel crosslinking. The stamps for hydrogel structuring were fabricated using a stereolithography printer (Formlabs Inc., Somerville, MA, USA) with CLEAR resin. Asiga® Flash Cure Box (405 nm, Asiga, Sydney, Australia) was used to cure the samples. Cell detachment was performed using Accutase (Thermo Fisher Scientific, Waltham, MA, USA). Fibronectin (Thermo Fisher Scientific, Waltham, MA, USA) was used for hydrogel surface coating at a concentration of 10 µg/mL in sterile phosphate-buffered saline (PBS). Confocal imaging was performed using a Zeiss LSM 780 confocal microscope (Carl Zeiss AG, Oberkochen, Germany).

### Fabrication of the Optical Protection Model

For the Optical Protection Model, cells were seeded at 10,000 cells per well in white 96-well plates with transparent bottoms and cultured in phenol-free medium. Cells were maintained under standard incubation conditions, and the culture medium was refreshed every two days. After four days of culture, cells reached an estimated confluency of approximately 70–80%. Prior to cell exposure, artificial epidermal layers were sterilized by ultraviolet irradiation for 40 min inside a tissue culture hood. Three types of artificial epidermal layers were used: a model without polydopamine nanoparticles; a model with a low nanoparticle concentration (Skin Tone 1, ST 1); and a model with a high nanoparticle concentration (Skin Tone 6, ST 6). These layers were assigned to seven experimental conditions, summarized in Table 1.

**Table 1:** Sample designations for the Optical Protection Model experiment.

| **Designation** | **UV-B exposure** | **Overlying layer** | **Purpose** |
| --- | --- | --- | --- |
| **Ctrl -** | - | none | Baseline cellular activity (negative control) |
| **Ctrl (ST6)** | - | ST6 (high PDA) | Tests whether the pigmented layer itself affects cell viability in the absence of UV |
| **Ctrl +** | - | none | Positive control for apoptosis |
| **UV** | 30 mJ/cm<sup>2</sup> | none | Unprotected UV-B exposure |
| **UV + PDMS** | 30 mJ/cm<sup>2</sup> | unpigmented PDMS | Tests optical contribution of the PDMS matrix |
| **UV + ST1** | 30 mJ/cm<sup>2</sup> | ST1 (low PDA) | Low-pigmentation photoprotection |
| **UV + ST6** | 30 mJ/cm<sup>2</sup> | ST6 (high PDA) | High-pigmentation photoprotection |

The layers were placed on the culture medium, and before irradiation, the culture medium was carefully removed from the wells so that the artificial epidermal layers were gently placed directly onto the cell monolayers according to the assigned experimental group. The irradiation setup was placed inside a biosafety cabinet. The distance between the lamp and the cell surface was fixed at 10 cm. To minimize stray irradiation and cross-well exposure, the sides of the 96- well plate were shielded with black barriers. Cells were exposed to a target UV-B dose of 30 mJ/cm², corresponding to an exposure time of 36 s. Control groups were handled identically but were not exposed to UV-B radiation. Immediately after irradiation, the artificial epidermal layers were removed, fresh culture medium was added, and the cells were returned to the incubator. Apoptosis was assessed 24 h after UV-B exposure. Caspase-Glo® 3/7 reagent was added to each well according to the manufacturer’s instructions. The plates were incubated for 60 min at room temperature, protected from light. Luminescence was measured using a plate reader. Staurosporine (1 µM) was used as a positive control for apoptosis induction and was added to designated wells 3–4 h prior to the Caspase-Glo® assay. The method is illustrated in Figure 2.

**Figure 2:**
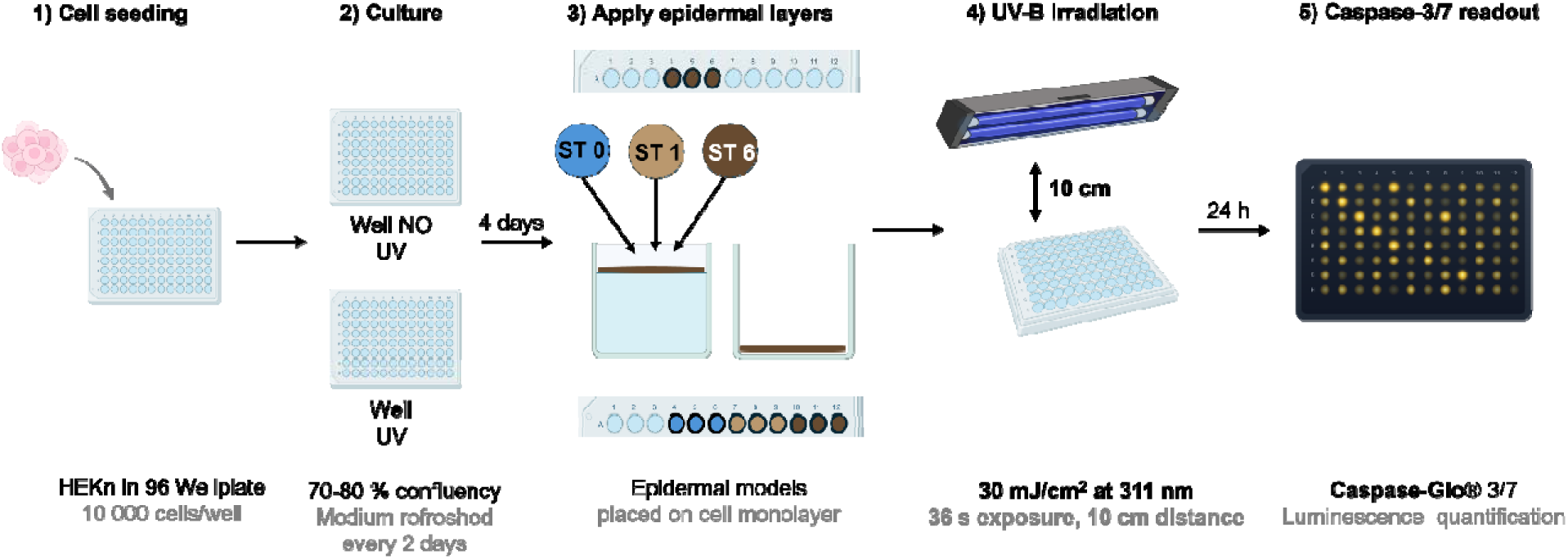
Schematic of the Optical Protection Model workflow: keratinocytes are seeded in 96-well plates, covered with artificial epidermal layers of varying pigmentation (low or high particle concentration), and exposed to UV-B irradiation. Apoptosis is quantified 24 h later using Caspase-Glo® 3/7 assay.

### Fabrication of the Structural Dermal Model

Hydrogel precursor solutions were prepared by dissolving two functional cfGel polymers, namely thiol-functionalized cfGel (cfGel-SH, 2.5 wt%) and norbornene-functionalized cfGel (cfGel-NB, 2.5 wt%) in PBS until fully dissolved to create a 5% precursor solution. For photo-crosslinking, an Irgacure 2959 photoinitiator (0.5%) solution was added to the hydrogel precursor solutions and mixed thoroughly to ensure uniform distribution. Structured hydrogel scaffolds were fabricated using a mold-based stamping approach. Stamps were designed and produced using a stereolithography 3D printer (Formlabs Inc., Somerville, MA, USA) with CLEAR resin (Figure 3B,C). Each stamp featured cylindrical protrusions with a length of 1.6 mm and a diameter of 0.4 mm, of which 0.8 mm was inserted into the hydrogel to define the resulting channel depth (Figure 3B). 300 microliters of hydrogel precursor solution were dispensed into the wells of tissue culture plates to reach a thickness of 1 mm. The plates were gently shaken to promote uniform filling of the wells and then left standing in the wells to allow trapped air bubbles to rise and dissipate. Subsequently, the stamps were carefully placed onto the hydrogel-filled wells. The hydrogel was crosslinked for 2 minutes. After crosslinking, the stamps were released by adding a small amount of PBS to the wells. The PBS reduced friction between the polymerized hydrogel surface and the stamp features, enabling gentle removal of the stamps without damaging the hydrogel structures. Prior to cell seeding, hydrogel scaffolds were coated with fibronectin to promote cell adhesion (200 µL, 10 µg/mL in sterile PBS). Wells were incubated at 37°C for 60 minutes, after which the fibronectin solution was carefully aspirated from the side of each well. Cells were expanded to 70–80% confluency prior to seeding. Cells were detached using 2 mL of pre-warmed accutase for 7 minutes at 37°C, and accutase activity was neutralized with 4 mL of pre-warmed medium. The suspension was centrifuged at 450 × g for 5 minutes, and the pellet was resuspended in fresh medium. Cells were seeded at 150 µL per well onto fibronectin-coated hydrogels, corresponding to approximately 6.28 × 10□ cells per well. Following seeding, the plate was transferred immediately to the incubator (37°C, 5% CO□) and left undisturbed for 4 hours for cell attachment. After this period, 350 µL of pre-warmed complete medium was added gently to each well. Medium was changed every other day. The method is illustrated in Figure 3A. Confocal images were taken 24 h and 5 days post-cell seeding.

**Figure 3:**
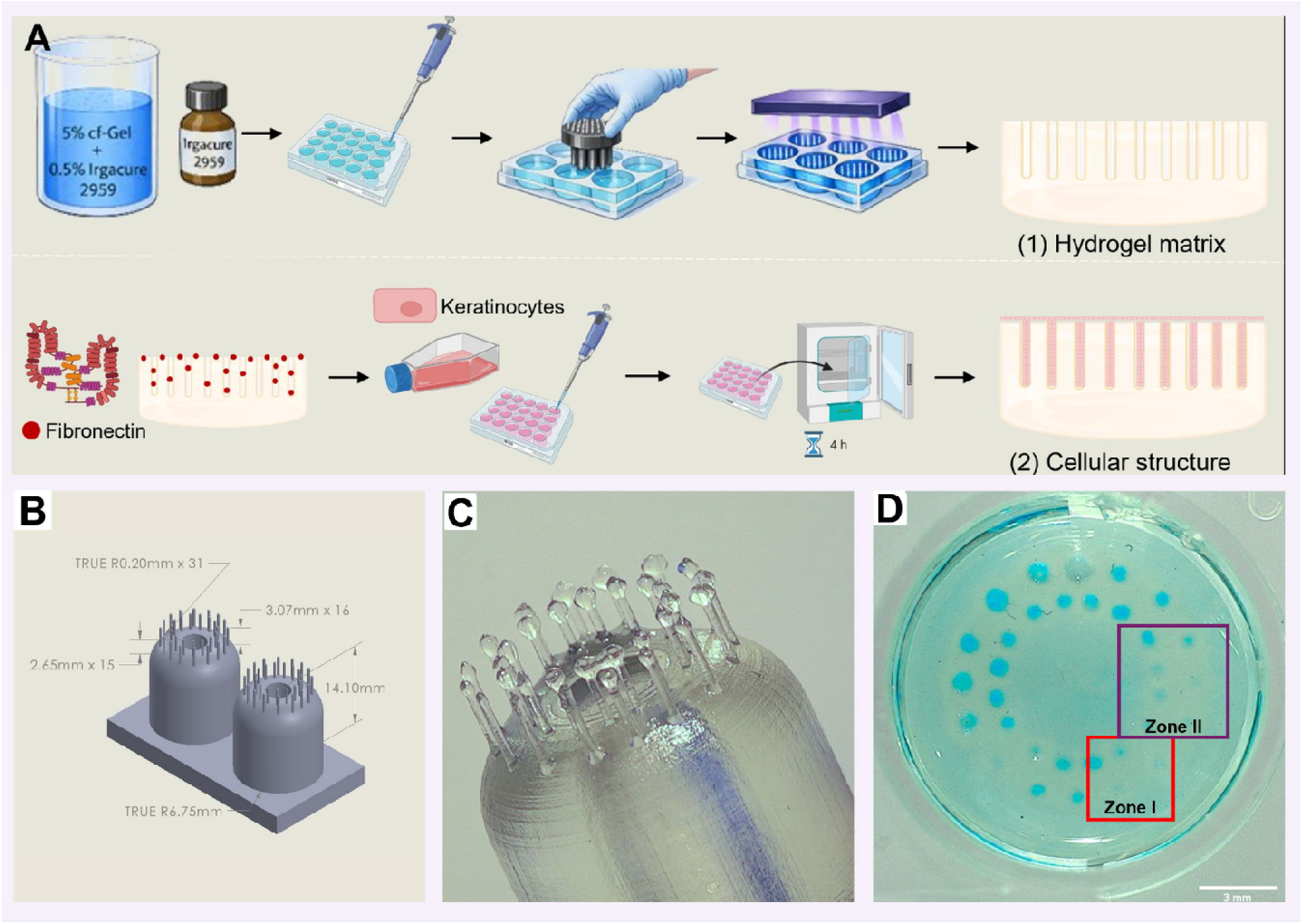
Fabrication of the Structured Dermal Model. (A) Schematic workflow of the two-step fabrication process. Top row: hydrogel matrix formation. Bottom row: cellular functionalization. (B) CAD design of the SLA-printed stamp featuring cylindrical protrusions. (C) Close-up photograph of the SLA stamp, illustrating the geometry and arrangement of the cylindrical protrusions (D) Macroscopic photograph of a structured cf-gel hydrogel scaffold showing the regular array of follicle-like channels after stamp removal. The red inset (zone I) and violet inset (zone II) mark the regions imaged in Figure 4B and Figure 4C–F, respectively. Scale bar: 3 mm.

## RESULTS AND DISCUSSION

### 3.1 The Optical Protection Model

The first demonstrator of the biohybrid model combines human keratinocytes with an optically defined artificial epidermis layer to study UVB-induced apoptosis under controlled optical conditions. A 30 mJ/cm² UVB dose was selected to give a robust yet sub-necrotic apoptotic response, consistent with values reported for primary keratinocytes in comparable *in vitro* settings ^12, 33, 34^. The results reveal a concentration-dependent relationship between the pigmentation of the artificial epidermal layer and the cellular apoptotic response (Figure 4A).

**Figure 4:**
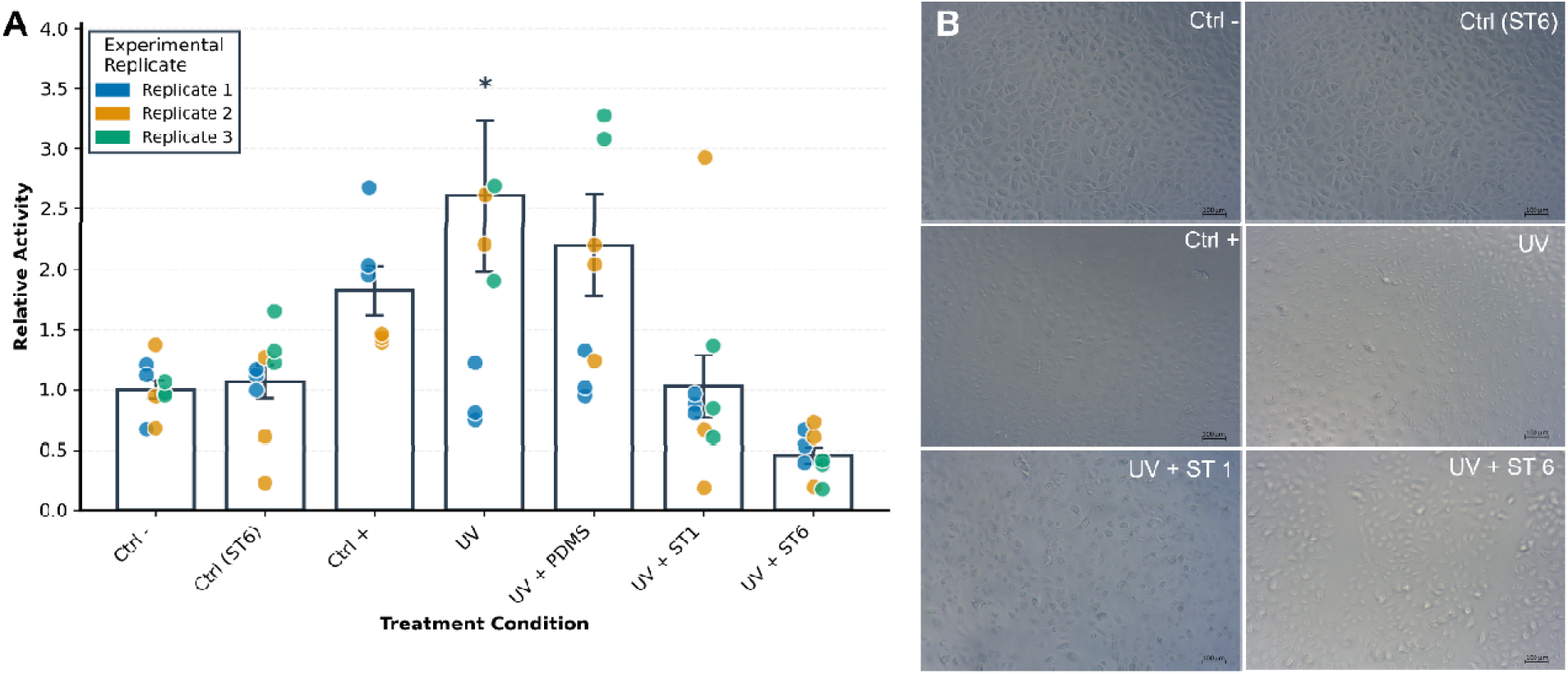
UV-B-induced apoptosis in keratinocytes under optically tunable artificial epidermal layers. **(A)** Relative caspase-3/7 activity across treatment conditions. Bars represent mean ± SEM; individual data points represent technical wells, color-coded by experimental replicate (n = 3 independent replicates per condition). **(B)** Representative brightfield images of keratinocytes under different conditions. Scale bar: 200 μm.

UVB exposure without optical protection (UV condition) induced significant apoptotic activity, with relative caspase-3/7 levels increasing 2.61 ± 0.63-fold (mean ± SEM, n = 60 across 3 independent replicates) compared to unexposed controls (Ctrl -; p = 0.014). This dose lies within the established range commonly used to model acute UV-B damage *in vitro*, where measurable caspase activation and DNA damage occur without causing extensive necrosis or complete loss of monolayer integrity ^12, 35^.

Crucially, the artificial epidermal layers increasingly attenuated this apoptotic response in proportion to their optical density. Models with low pigmentation (UV+ST1) reduced relative caspase-3/7 activity to 1.03 ± 0.26, which was significantly lower than the unprotected UV condition (p = 0.017) and indistinguishable from the unexposed Ctrl− group at the level of pairwise testing (p = 1.000). High-pigmentation models (UV + ST6) provided substantially stronger protection, reducing activity to 0.46 ± 0.07, significantly below both the unprotected UV condition (p = 0.0003) and the UV + PDMS condition (p = 0.006). In contrast, the addition of an unloaded PDMS layer (UV + PDMS, mean = 2.20 ± 0.42) did not significantly reduce apoptosis relative to the unprotected UV condition (p = 0.972), confirming that the PDMS matrix itself contributes negligibly to optical attenuation at 311 nm and that the photoprotective effect is attributable exclusively to the PDA nanoparticle content. The Ctrl (ST6) condition (mean = 1.07 ± 0.14, p = 1.000 vs. Ctrl−) confirmed that contact with the pigmented artificial epidermal layer does not in itself induce apoptosis under the experimental conditions used. Notably, the UV+ ST6 mean of 0.46 fell below the unexposed control (Ctrl-). Several factors may contribute to this observation. The most plausible is a cell loss artifact during model removal in the form of mechanical detachment of the ST6 layer that may lift weakly adherent cells, resulting in fewer cells remaining in the well at the time of the measurement. Another explanation could be that necrosis, rather than apoptosis, which potentially would be induced by mechanical stress during removal, would not register in the caspase-3/7 readout and could additionally suppress the signal. The relative contributions of these mechanisms cannot be resolved from the current dataset. This artifact does not, however, undermine the core conclusion of the experiment, that increasing pigmentation produces a stepwise reduction in UV-induced apoptotic signaling. The monotonic reduction in caspase activity with increasing pigmentation (UV > UV + ST1 > UV + ST6) is preserved across all replicates, and the intermediate value observed for UV + ST1, which falls between the unprotected UV condition and UV + ST6, cannot be explained by mechanical artifact alone and is consistent only with a genuine, pigmentation-dependent optical effect. This quantitative protection correlates remarkably well with the measured optical transmission of the artificial epidermal layers at 311 nm. Skin Tone 1 transmits ∼75% of incident UV-B, while Skin Tone 6 transmits ∼10% ^23^. The near-linear relationship between optical attenuation and biological response validates the fundamental assumption of this model. The positive control (Ctrl +) showed a mean relative activity of 1.83 ± 0.20. However, this condition was based on only two independent experimental replicates, limiting statistical power, and the comparison to Ctrl- did not reach significance (p = 0.670). Staurosporine values are therefore reported for transparency but are not used to draw biological conclusions.

Brightfield microscopy provided qualitative confirmation of the quantitative caspase data (Figure 4B). The control cells exhibited typical keratinocyte morphology, a polygonal shape with clear cell boundaries, indicating intact cell-substrate adhesion. UV-exposed cells without protection showed characteristic apoptotic morphology. UV-exposed cells without protection showed marked cell rounding and reduced substrate adhesion, morphological features consistent with apoptosis ^34, 36^, and comparable to the positive control group. These features were reduced in cells protected by highly pigmented models, resulting in a morphology more similar to that of unexposed controls. However, signs of apoptosis are evident in both the ST 1 and ST 6 samples. It is important to acknowledge the simplifications in this model. The 2D monolayer geometry eliminates the structural complexity of stratified epidermis, including the three-dimensional architecture of rete ridges ^37, 38^. This simplification isolates the purely optical component of photoprotection and is appropriate for systematic device testing or material characterization, but direct extrapolation to *in vivo* photoprotection may require consideration of additional factors, including tissue architecture or continuous vs. acute exposure. The unprotected UV condition exhibited higher inter-replicate variability (SEM = 0.63, coefficient of variation across day-means ≈ 53%) than the protected conditions. In particular, one of three replicates showed a substantially attenuated caspase response in the unprotected UV group, which we attribute to the inherent sensitivity of primary keratinocyte UV-B response to subtle differences in cell handling, seeding density, and lamp warm-up state. This variability should be considered when interpreting pairwise comparisons involving the UV group. Similarly, apoptosis represents only one of several UV-B responses. Keratinocytes also activate DNA repair mechanisms, inflammatory signaling, and oxidative stress responses that occur on different timescales ^25, 39^ and should be considered in future studies.

The Optical Protection Model could enable various applications where systematic optical control is advantageous. These include sunscreen testing under standardized irradiation conditions ^33, 40, 41^, wavelength-specific photodamage studies by substituting different light sources, and calibration of optical sensors for dermatological applications. Additionally, the 96-well plate format further enables high-throughput screening, which is important in industrial settings.

### 3.2 Formation of Cell-Seeded Follicle-Like 3D Structures

The Structured Dermal Model extends the biohybrid model beyond planar geometry and show that optically defined structures can be combined with living cells in three dimensions. Hydrogel matrices made from cold-water fish gelatin (cf-Gel) were chosen as the scaffold material due to their established biocompatibility, adjustable mechanical properties, and suitability for photocrosslinking ^32^. Importantly, cf-Gel remains liquid at room temperature, which enables easy handling during the crosslinking phase ^32^. Hydrogel scaffolds with arrays of follicle-like microcavities were fabricated using the stamping protocol described in Methods (Figure 3A–D).

The cylindrical geometry was selected as a simplified representation of the elongated morphology of hair follicles while remaining compatible with the limitations of the fabrication methods. Although native hair follicles exhibit a more complex tapered structure and contain multiple concentric tissue layers, the cylindrical cavities capture the essential feature of a confined vertical microenvironment in which cells can accumulate and organize. The microcavity diameter of 400 µm is larger than the diameter of human hair shafts (∼50–100 µm)^42, 43^, but within the range of follicle opening dimensions, the depth of 800 µm is approximately one quarter of native human follicle length and was selected to remain within the working distance of standard confocal objectives. Figure 3D shows a macroscopic top-down view of the resulting construct within a standard culture dish. Macroscopic inspection of the stained hydrogel array shows generally regular spacing of the microcavities, though local variation in channel diameter and depth was observed across the scaffold. This variability likely comes from minor inconsistencies in stamp-hydrogel contact pressure during crosslinking and removal. The variabilities show the sensitivity of the fabrication process to mechanical alignment and pressure distribution, suggesting that future iterations could benefit from controlled compression setups to improve the models.

The modular assembly of the biohybrid construct is shown in Figure 5A in both top-down and lateral views. The artificial epidermal layer sits in conformal contact with the hydrogel surface, with no visible gap at the interface, confirming that the stacked geometry allows straightforward exchange of the optical top layer without disturbing the underlying cellular scaffold. Figure 5B shows a brightfield micrograph of the hydrogel surface within the channel array region (corresponding to the red inset in Figure 3D, zone I). Brightfield microscopy at 24 h post-seeding revealed that keratinocytes did not distribute uniformly across the hydrogel surface. Instead, cells were predominantly found as compact multicellular aggregates concentrated in the microcavities, while the regions between cavities showed comparatively sparse coverage. This pattern suggests that passive gravitational settling directed cells preferentially toward the channel openings, where individual cells accumulated and subsequently formed compact multicellular clusters during the 4-hour attachment period ^44, 45^. Whether cells penetrated the full channel depth (0.8 mm) or accumulated primarily at the entrance could not be resolved from surface brightfield imaging.

**Figure 5:**
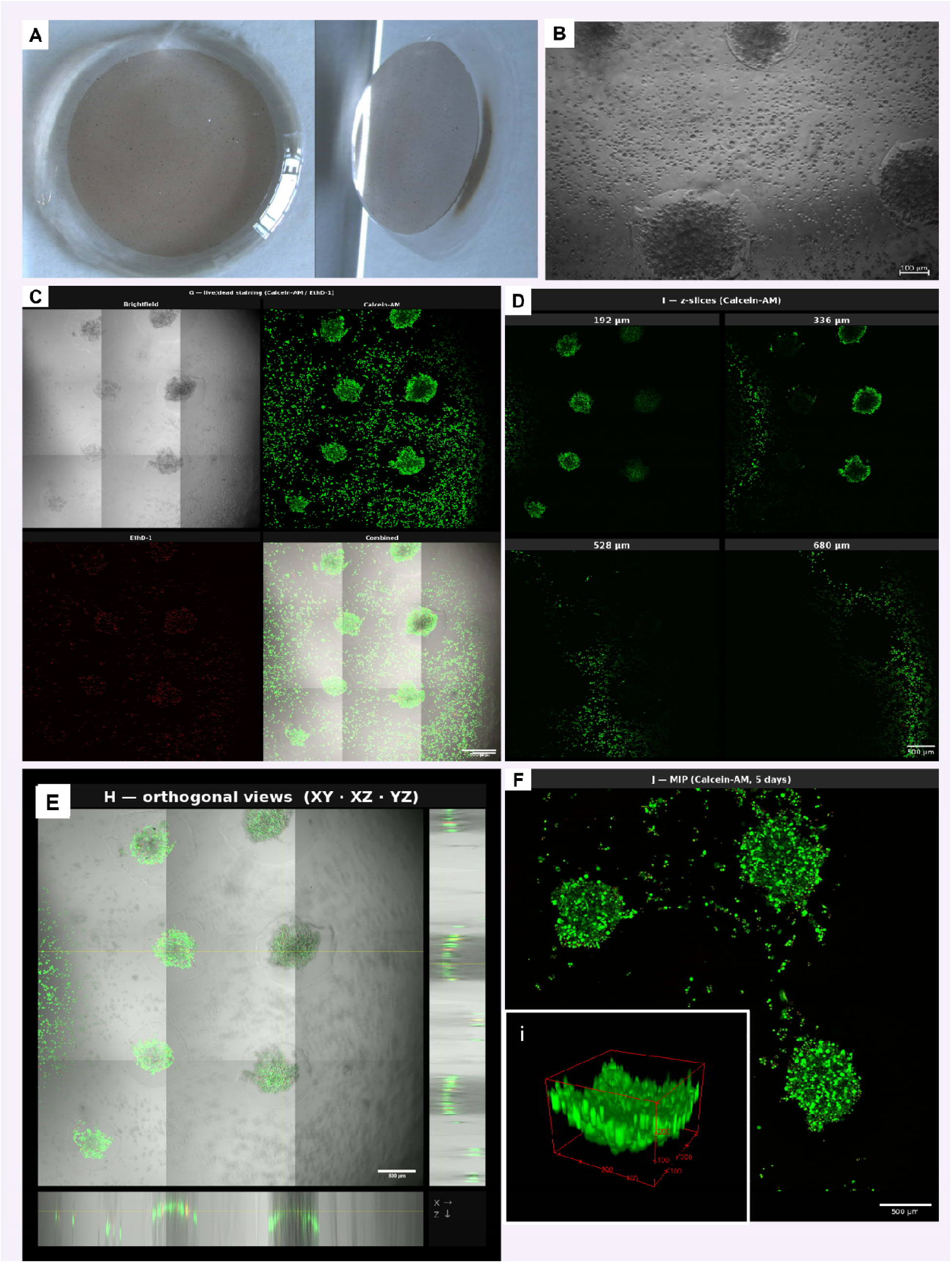
Cellular characterization of the Structured Dermal Model. (A) Top-down and lateral views of the assembled biohybrid model. (B) Brightfield micrograph of HEKn seeded on fibronectin-coated hydrogel scaffolds at 24 h post-seeding. Scale bar: 100 µm. (C) Confocal fluorescence images of live/dead-stained constructs at 24 h post-seeding. Top row: brightfield and Calcein-AM channel (green, viable cells). Bottom row: EthD-1 channel (red, membrane-compromised cells) and combined overlay. Scale bar: 500 µm. (D) Representative z-slices of the Calcein-AM channel at depths of 192, 336, 528, and 680 µm, showing the vertical distribution of viable cells. Scale bar: 300 µm. (E) Orthogonal views (XY, XZ, YZ) of the combined brightfield and Calcein-AM z-stack. Scale bar: 500 µm(F) Maximum intensity projection (MIP) of the Calcein-AM channel at 5 days post-seeding. Inset (i): three-dimensional surface rendering of a representative microcavity. Scale bar (main panel): 500 µm.

Cell viability within the three-dimensional constructs was assessed by Calcein-AM/EthD-1 live/dead fluorescence staining, imaged by confocal microscopy (Figure 5C, corresponding to zone II in Figure 3D). In the Calcein-AM channel, strong green fluorescence is detected in compact multicellular clusters localized within and adjacent to the microcavities, confirming robust metabolic activity at 24 hours post-seeding. In the EthD-1 channel, only a few scattered red signals are visible, indicating that most cells remained membrane-intact at this time point. The spatial concordance between the brightfield morphology and Calcein-AM signal in the overlay panels confirms that the fluorescent clusters correspond to viable keratinocytes. To analyze the spatial distribution of viable cells across the full depth of the scaffold, representative z-slices were shown at four different axial positions: 192, 336, 528, and 680 µm (Figure 5D). At shallow depth (192 µm), the cells are detected in multicellular clusters situated at the bottom of the cavities. The 192 µm z-axis section corresponds to a deeper focal plane that captures the signal from within the follicle-like microcavities where the cell clusters have accumulated. At this depth, green fluorescence is observed in clearly defined multicellular clusters that correspond to the location of the cavities, confirming that the cells remain viable within the channel structures. At 336 µm depth, the cells in the outer ring are visible through green fluorescence. At shallower depths (528–680 µm), the fluorescence signal increasingly spreads across the surface of the scaffold, consistent with the presence of cells on the surface of the hydrogel rather than within the channels. The spatial pattern observed along the z-positions is most likely due to an irregular topography of the hydrogel surface within the culture well. During crosslinking, the hydrogel precursor solution tends to accumulate in greater thickness along the walls of the well compared to the center, resulting in a meniscus-like geometry. This means that the absolute depth at which the openings of the microcavities are located varies across the scaffold, with central regions having channel openings at shallower absolute z-positions and peripheral regions at greater depths. Therefore, the apparent change in cluster localization along the z-sections reflects the curved scaffold geometry rather than a true vertical redistribution of cells.

The three-dimensional architecture of cell distribution is illustrated by orthogonal projections of the combined brightfield and Calcein-AM z-stack (Figure 5E). The XZ and YZ cross-sectional views reveal that viable cells are not distributed homogeneously throughout the channel volume but instead form a curved, arc-like layer within the lower portion of the microcavities. This curvature reflects the cells conforming to the cylindrical geometry of the channel walls rather than settling as a flat horizontal layer (Figure 5F, inset i). The upper portions of the channels remain largely unpopulated. The observations indicate that while the microcavities successfully capture and confine cells at their base, a full three-dimensional population of the channel walls is not achieved under the current passive seeding conditions. To evaluate whether keratinocyte viability is maintained beyond the initial attachment phase, constructs were additionally imaged at five days post-seeding and illustrated using a maximum intensity projection (MIP) (Figure 5F). At this timepoint, Calcein-AM signal remains detectable within the microcavity positions, confirming that keratinocytes retain metabolic activity over a culture period of five days. This observation provides early evidence that the dermal model supports not only initial cell adhesion but also sustained viability, which is a basic prerequisite for the model’s use in functional endpoint assays beyond acute readouts. Whether cells are actively proliferating within the channels or simply maintaining viability at stable numbers was not determined.

In summary, this demonstrator proves the feasibility of combining structured hydrogel scaffolds, living keratinocytes, and tunable artificial epidermal layers within a single modular model. The application shows how this decoupled architecture, with a top layer with defined optical properties and an interchangeable bottom hydrogel scaffold with cells, represents a new testing principle absent from existing tissue-engineered or synthetic skin models. The z-slice series and orthogonal views confirm three-dimensional cell localization within the microcavities, while the five-day culture data demonstrate that the model supports sustained cellular viability beyond the initial attachment phase. While the present implementation uses HEKn as a model cell type, the approach is fundamentally transferable to other cell types, including dermal papilla cells or melanocytes ^44^. Additionally, the interchangeable architecture of the dermal part enables variable modulation according to the specific need. The gelatin hydrogel provides a matrix that supports the adhesion and culture of multiple skin-relevant cell types ^32^, and the fibronectin coating used here is a generic adhesion ligand rather than a keratinocyte-specific factor. The scaffold geometry and crosslinking chemistry are therefore independent of the cell type introduced, enabling straightforward substitution or co-seeding without modification of the fabrication workflow.

## Conclusion

This paper presents a modular biohybrid skin model that integrates an optically defined artificial epidermal model with living keratinocytes. The two demonstrators, the Optical Protection Model and the Structured Dermal Model, illustrate the model’s fundamental principle: a controlled optical model can coexist with biologically reactive cellular components within a single, reconfigurable test system.

The Optical Protection Model confirmed the model’s fundamental assumption by revealing a quantitative, linear relationship between the pigmentation of the skin model and the UV-induced apoptotic response. This dose-dependent relationship, which was reproduced in three independent replicate experiments, confirms that the optical properties of the artificial epidermal model layers can be predictably translated into measurable biological outcomes at the cellular level. The Structured Dermal Model extends the principle to a geometrically structured hydrogel scaffold and shows that keratinocytes remain viable in follicle-like microcavities within this construct. Taken together, the results show the first steps toward a new class of optical skin model, while also identifying the development steps required before it can be applied to specific testing scenarios. At the biological level, the present study focused on acute endpoints such as short-term viability and UV-induced apoptosis, for which the current model provides biologically meaningful readouts. Extended culture periods, proliferation assays, and functional outcomes would address complementary questions and should be assessed in future work. Beyond the two demonstrations presented here, the modular architecture of the bio-hybrid model opens a range of concrete testing applications that combine optical control with cellular readout. By applying topical formulations to the pigmented artificial epidermal layer and quantifying the resulting reduction in UV-induced caspase activity of the underlying keratinocyte monolayer, the model could enable sun protection factor (SPF)-style measurements under controlled, skin-tone-specific optical conditions. The same architecture could be adapted to further photomedicine workflows, including intense pulsed light (IPL) and laser-based hair removal on the Structured Dermal Model ^46, 47^, as well as photodynamic and photothermal therapy (PDT/PTT) of skin malignancies, with light-induced cell death quantified as a function of overlying pigmentation and wavelength. The platform could further support the validation of optical medical devices and wearable sensors such as pulse oximeters and photoacoustic imagers, where performance disparities across skin tones have been extensively documented but rarely addressed at the level of standardized testing ^48–50^. Demonstrating these use cases quantitatively and benchmarking the model against established alternatives is the natural next step. This model represents a novel class of bio-hybrid skin model and, in accordance with the 3R principles, has the potential to reduce the use of animal models and primary human tissue in the early development phase of optical devices.

By combining the reproducibility and optical tunability of synthetic models with the biological responsiveness of living cells, the model represents a novel approach between inert optical test systems and variable tissue-engineered models, which shows great promise as optical biomedical technologies increasingly require both physical precision and biological relevance in their validation processes.

## Acknowledgments

This research received no specific funding. The authors used Grammarly and PaperPal for language editing and grammar correction.

